# Cerebellar disruption impairs working memory during evidence accumulation

**DOI:** 10.1101/521849

**Authors:** Ben Deverett, Mikhail Kislin, David W. Tank, Samuel S.-H. Wang

**Affiliations:** Princeton Neuroscience Institute, Princeton University, Princeton, NJ 08544, USA; Department of Molecular Biology, Princeton University, Princeton, NJ 08544, USA; Rutgers Robert Wood Johnson Medical School, Piscataway, NJ 08854, USA

## Abstract

To select actions based on sensory evidence, animals must create and manipulate representations of stimulus information in memory. We found that during accumulation of somatosensory evidence, optogenetic manipulation of cerebellar Purkinje cells reduced the accuracy of subsequent memory-guided decisions and caused mice to downweight prior information. Behavioral deficits were consistent with the addition of noise and leak to the evidence accumulation process, suggesting the cerebellum can influence the maintenance of working memory contents.

---

The accumulation of sensory evidence is an important part of decision-making^1^. In rodents performing evidence accumulation, neuronal perturbation of specific brain regions can have distinct effects on behavior^2^. Depending on the region, perturbation can cause minimal effects^3^, it can impair functions related to decision-making^3–6^, or it can influence evidence integration in working memory^7^. Many forebrain regions implicated in evidence accumulation receive input from the lateral posterior cerebellum^8–10^, and disruption of the human cerebellum produces working memory impairments^11–14^. Given its roles in sensorimotor integration^15^ and motor preparation^16^, cerebellar output may influence the evidence accumulation process. Here we examined whether direct, temporally precise disruption of cerebellar neural activity modulates the accumulation of somatosensory evidence.

We used a behavioral task for head-fixed mice in which animals accumulate sensory evidence over a period of seconds to guide decisions^17^. In each trial (Fig. 1a) the mouse is presented with simultaneous streams of randomly timed left- and right-sided whisker puffs followed by a delay, after which it licks in the direction of more puffs to retrieve a water reward. We previously showed that coarse full-session pharmacological perturbation of the lateral posterior cerebellum alters performance in this task, and that Purkinje cell (PC) activity there encodes stimulus- and decision-related variables^17^. In the present study we trained 13 mice on this task over hundreds of behavioral sessions (Fig. 1b, Supplementary Fig. 1).

**Figure 1:**
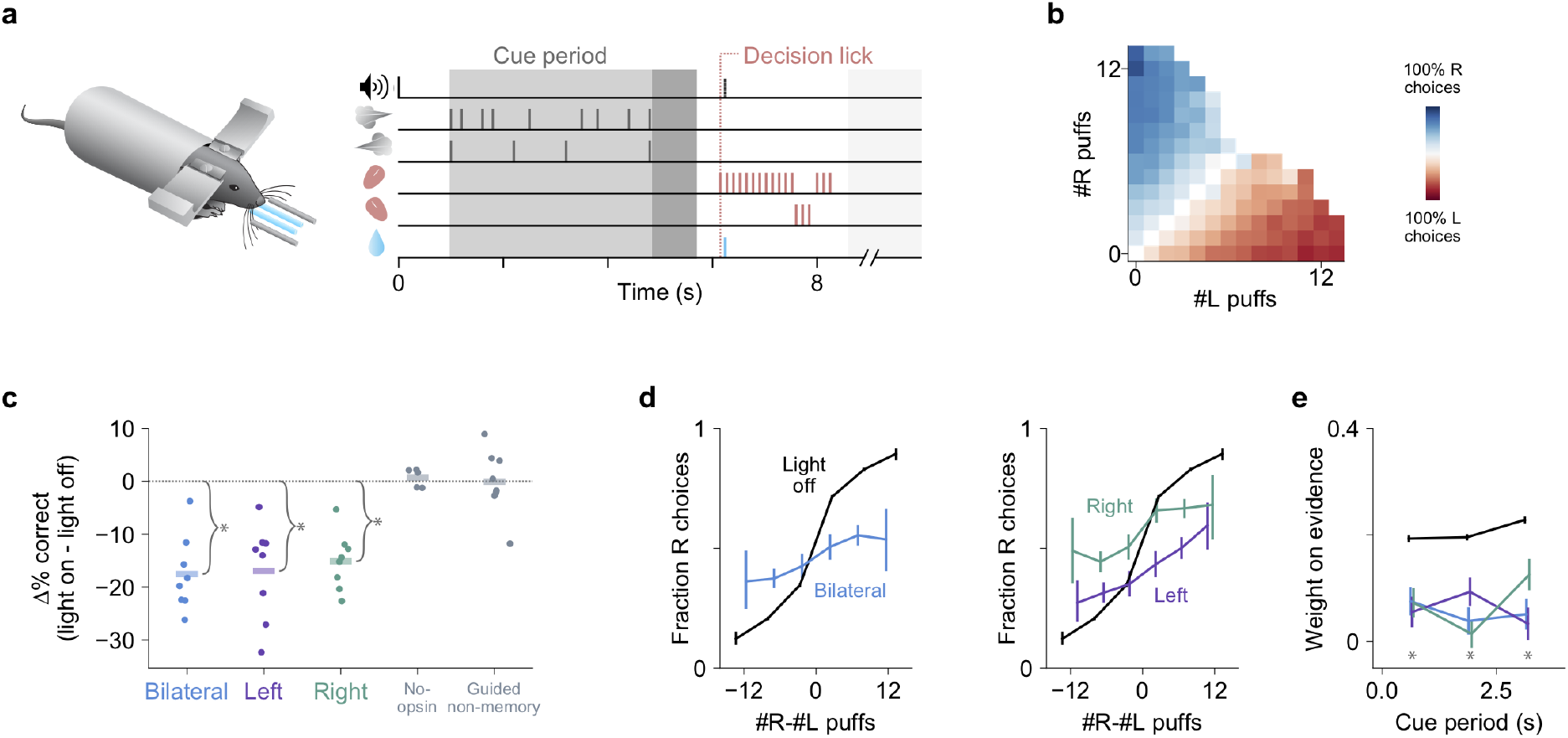
Cerebellar disruption during evidence accumulation impairs decisions. (a) Schematic of the evidence-accumulation decision-making task. In each trial, two streams of random, temporally Poisson-distributed air puffs were delivered to the left and right whiskers. After an 800-ms delay, mice licked one of two lick ports indicating the side with more cumulative puffs to receive a water reward. Gray-shaded regions from left to right: cue period, delay, intertrial interval. Decision lick: first detected lick after the delay. (b) Choice probabilities as a function of the number of left- and right-side puffs (n=96,254 trials over 664 sessions in 13 mice). (c) Change in performance as a result of cue-period light delivery to the left, right, or bilateral cerebellum (n=46,435 light-off trials, 5,392 light-on trials, 397 sessions, 8 mice). Dots: individual mice. Lines: mean across mice. *: p<0.01 (two-tailed paired t-test). No-opsin: bilateral light delivery in ChR2^−^ mice (also see Supplementary Fig. 3). Guided non-memory: bilateral light delivery in trials where mice were guided to lick the correct side by delivery of all-single-sided puffs during the cue period and delay. (d) Psychometric curves for light-off (black) trials and light-on (colored) trials from all perturbation sessions in all experimental mice. Results are shown for bilateral (left) and unilateral (right) perturbations. Error bars: 95% CI. (e) Regression of animal choices on evidence quantity throughout the cue period for light-off (black) and light-on (colored) trials. Weights indicate the extent to which evidence was used to guide decisions, and the sum of weights is proportional to overall performance. *: p<0.01 (99% CI, light-off: 0.18–0.21, 0.18–0.21, 0.21–0.25; bilateral: 0.01–0.15, -0.03–0.11, -0.02–0.13; left: -0.02–0.13, 0.02–0.16, -0.04–0.11; right: 0–0.14, -0.05–0.08, 0.05–0.2)

To determine whether cerebellar activity can modulate the evidence accumulation process, we used time-resolved, cell-type-specific optogenetic perturbation specifically during the cue period, when evidence is presented and prior to the decision. We stimulated ChR2-expressing PCs (Supplementary Fig. 2), which inhibit the cerebellar output nuclei, using light delivered through optical fibers implanted bilaterally over crus I of the cerebellum. Light was delivered for the full duration of the cue period, either bilaterally or unilaterally in a randomly selected subset (15-30%) of trials over hundreds of behavioral sessions in 8 ChR2-expressing mice. Both unilateral and bilateral cerebellar perturbations led to reductions in performance (Fig. 1c-e), and unilateral perturbation induced a small ipsilateral choice bias on average (Fig. 1d). Impaired performance was associated with downweighting of evidence throughout the cue period (Fig. 1e). As a negative control, light delivery did not alter performance in ChR2^−^ mice (Fig. 1c, No-opsin; Supplementary Fig. 3).

In this experiment, the decision lick occurred approximately 1 second (1.31 ± 0.29 s, mean ± s.d.) after the end of light delivery, suggesting that the impairment did not arise from a deficit in the ability to lick. We nevertheless considered that light delivery might introduce a delayed effect that interfered with motor readout. Three measurements suggest otherwise. First, the fraction of trials in which animals made a response (in either direction) was unaffected by the perturbation (98.6 ± 1.8% mean ± s.d. in light-on trials vs 99.7 ± 0.3% in light-off trials; p=0.11, two-tailed paired t-test). Second, the latency from the end of the delay period to the decision lick was indistinguishable between light-on and light-off trials (578 ± 222 ms mean ± s.d. light-off vs 595 ± 332 ms light-on; p=0.19 bilateral, p=0.84 left, p=0.14 right, two-tailed paired t-test within subjects). Finally, light delivery did not influence the ability to make directed decision licks in trials where mice were cued which direction to lick with all-unilateral puffs during the cue period and delay (Fig. 1c, Guided non-memory; Supplementary Fig. 3). Therefore, cerebellar disruption during the cue period affected not the ability to lick but rather one or more aspects of the preceding process.

The observed impairment could be explained by a variety of mechanisms, including alteration of the weight of incoming stimuli (i.e. sensory gating or attention), impairment of the retention of past stimulus information, or interference with translation of accumulated information into directed motor actions^16^ (Supplementary Fig. 4). We tested these alternatives by introducing additional trials in which light was delivered during a subsection of the cue period (Fig. 2). By regressing animal choice on evidence strength throughout the cue period (as in Fig. 1e), we quantified which specific cues animals remembered and incorporated into their choices, lending insight into the contents of their working memory when light was applied. Importantly, this approach differentiates scenarios that appear similar with simpler analyses, such as one in which light resets the animal’s retention of accumulated evidence vs. one in which accumulation is intact but light prevents the animal from executing the desired lick (Supplementary Fig. 4).

**Figure 2:**
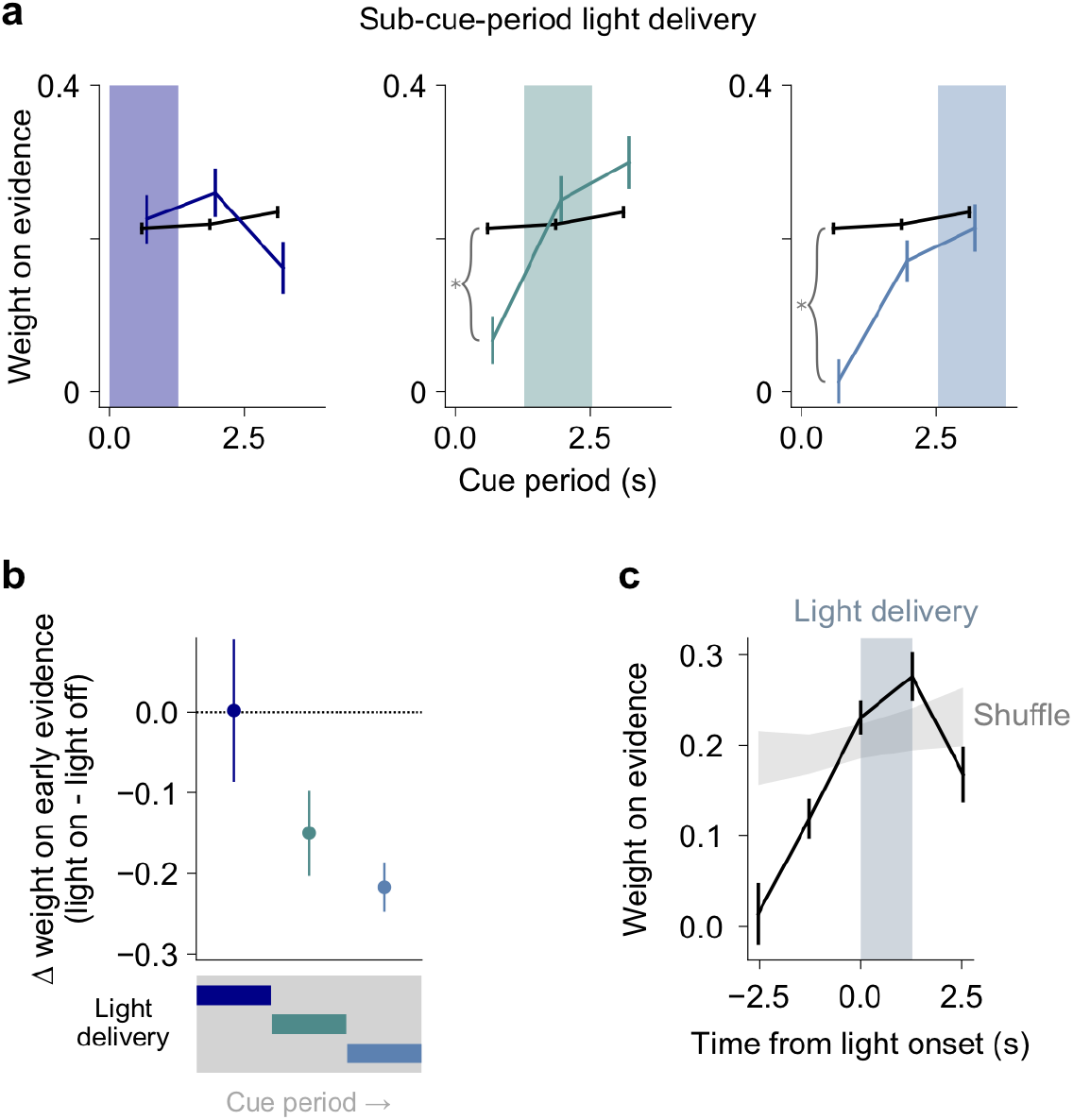
Cerebellar disruption influences weighting of past evidence. (a) Regression of animal choices on evidence quantity for light-off (black) and light-on (colored) trials (n=32,311 light-off trials, 5669 light-on trials, 285 sessions, 8 mice). Weights indicate the extent to which evidence was used to guide decisions, and the sum of weights is proportional to overall performance. Colored shading indicates the time of light delivery. Error bars: s.e.m. of regression weights. *: p<0.01 (99% CI on first bin, light-off: 0.19–0.23; light-on middle third: -0.01–0.15; light-on last third: -0.06–0.09). (b) Change in weight on evidence in the first third of cue period, as a function of when light was delivered during the cue period. Data points and error bars show mean ± s.e.m. across mice. (c) Evidence weight as a function of time relative to the onset of light delivery, with all cue-period light delivery conditions included (see Methods). Shuffle: light delivery time labels were shuffled before regression. Error bars: bootstrap s.d.

Surprisingly, mice had no difficulty using the evidence presented concurrent with light delivery, but they did have difficulty retaining evidence that had been previously presented (Fig. 2a-c). In the most extreme case, light delivery in the final third caused mice to completely discount evidence from the first third of the cue period (Fig. 2a right panel, first weight 95% CI: -0.04–0.07). In other words, light delivery in the middle and final third did not cause uniform effects across all trials, but instead selectively altered behavior in those trials where evidence was strong near the *start* of the cue period, prior to light delivery. In additional separate trials with light delivery during the post-evidence delay period, mice downweighted evidence throughout the entire preceding cue period (Supplementary Fig. 5).

These results suggest that cerebellar perturbation influenced behavior by altering how mice integrate and retain evidence information over time. We further tested this hypothesis by fitting our data to an established drift-diffusion framework that explicitly models the incremental integration of pulses of evidence to form decisions^18^. Crucially, this model differentiates impairments in evidence integration and storage per se (*e.g*. leakiness of evidence from memory) from non-specific impairments such as decision lapses that occur when animals fail to translate accumulated information into the proper action (Supplementary Movie 1, Supplementary Fig. 6). The model achieves specificity by taking advantage of the broad statistical distribution of stimulus timings available from thousands of trials.

Our model estimated parameters quantifying accumulator noise (σ_*a*_), sensory noise (σ_*s*_), memory leak or instability (λ), left-right bias, and a lapse rate. We fit all trials pooled across mice for the baseline light-off condition (n=56,550 trials), full-cue-period light delivery (n=6,394 trials), and delay-period light delivery (n=2,369 trials). Fits to light-off trials (Fig. 3a, top row, Supplementary Table 1) demonstrate that at baseline mice performed evidence accumulation using strategies similar to humans and rats^18^, with small values for accumulator diffusion noise and lapse rate, and with leaky accumulation (<0) consistent with the regression analysis (Supplementary Fig. 1b). When light was delivered for the full cue period (Fig. 3a, second row, Supplementary Table 1), behavior was characterized by an increase in *σ*_a_^2^, the diffusion noise in the accumulation process, and a decrease in λ, indicative of leakiness in evidence integration. Strikingly, the decay time constant τ (=1/λ) of accumulated evidence in working memory decreased approximately tenfold, from 6.7 s in the baseline condition to 0.72 s with light delivery. Therefore, cerebellar disruption impaired the noise and stability of accumulated working memory contents (Fig. 3b, Supplementary Movie 1). In contrast, when cerebellar activity was perturbed during the delay (Fig. 3a, bottom row, Supplementary Table 1), performance deficits were likely best explained by an increase lapse rate, consistent with disruptions to accumulated information or to translation of that information into actions.

**Figure 3:**
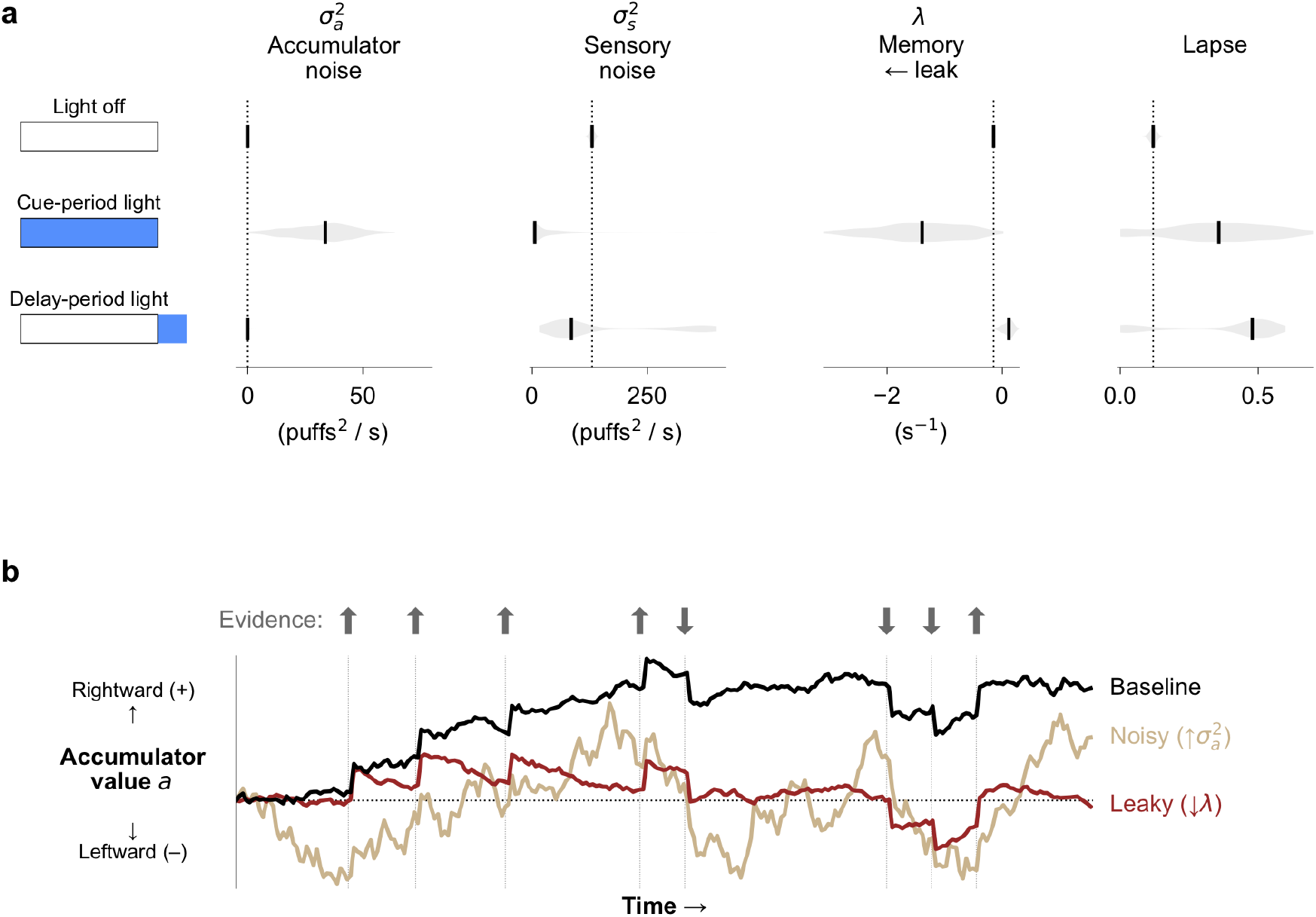
Fits to a drift-diffusion model reveal specific deficits in evidence accumulation. (a) Best-fit drift diffusion model parameters in different light delivery conditions (schematics on left indicate light delivery condition, with the box denoting the cue period and blue shading denoting light delivery). Fits were computed multiple times for each condition using random subsets of the data to assess the reliability of the best-fit parameters (see Methods). Black vertical ticks indicate the median best-fit parameter across fit repetitions. Gray shading represents the distribution of fit parameters across repetitions. Vertical dotted lines denote best-fit values in the light-off condition. (b) Visualization of the drift-diffusion model. The model’s accumulator value *a* is shown as it evolves over time in a single behavioral trial. Colored lines demonstrate how the trajectory of *a* is qualitatively altered by changes in specific parameters. Arrows and associated vertical lines indicate pulses of evidence. See also Supplementary Movie 1.

These results are consistent with clinical memory impairments observed after cerebellar lesions^11,12^, cerebellar roles in sensorimotor integration^15,19^, and theories of cerebellar function in working memory^20^. The results also align with recently reported cerebellar roles in motor preparation^16,21^, but add to those findings by extending cerebellar influence to the domain of evidence storage and manipulation for decision formation. The behavioral effects we characterized here were not observed with perturbations of other brain regions in similar paradigms^3–5,7^.

---

**Supplementary Figure 1:**
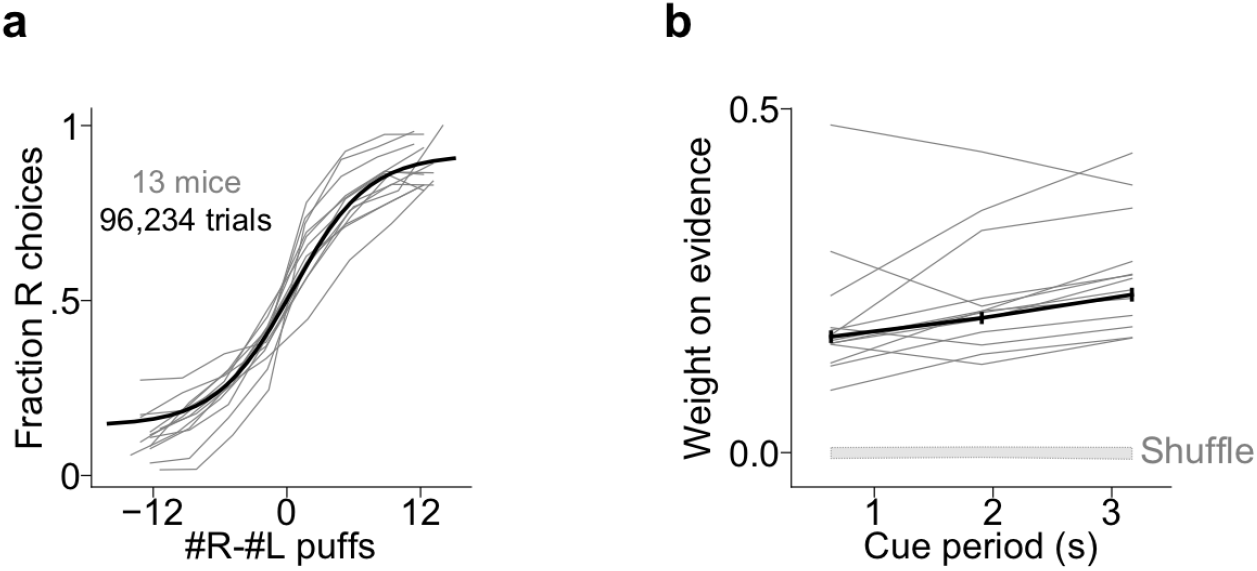
Performance in the somatosensory evidence accumulation task. (a) Psychometric curves for all individual mice (gray lines) and psychometric fit to the meta-mouse (black) consisting of all trials from all mice (n=96,254 trials over 664 sessions in 13 mice). Error bars: 95% CI. (b) Regression analysis demonstrating the extent to which mice use evidence throughout the cue period to guide decisions. Weights indicate the extent to which evidence was used to guide decisions, and the sum of weights is proportional to overall performance. Upward slope indicates a slight tendency to weight later evidence more heavily than earlier evidence (error bars show 95% CI), which would be predicted by leaky integration of stimuli. Gray lines: individual mice. Black line: meta-mouse. Bottom gray shading: 95% CI when choice was shuffled across trials.

**Supplementary Figure 2:**
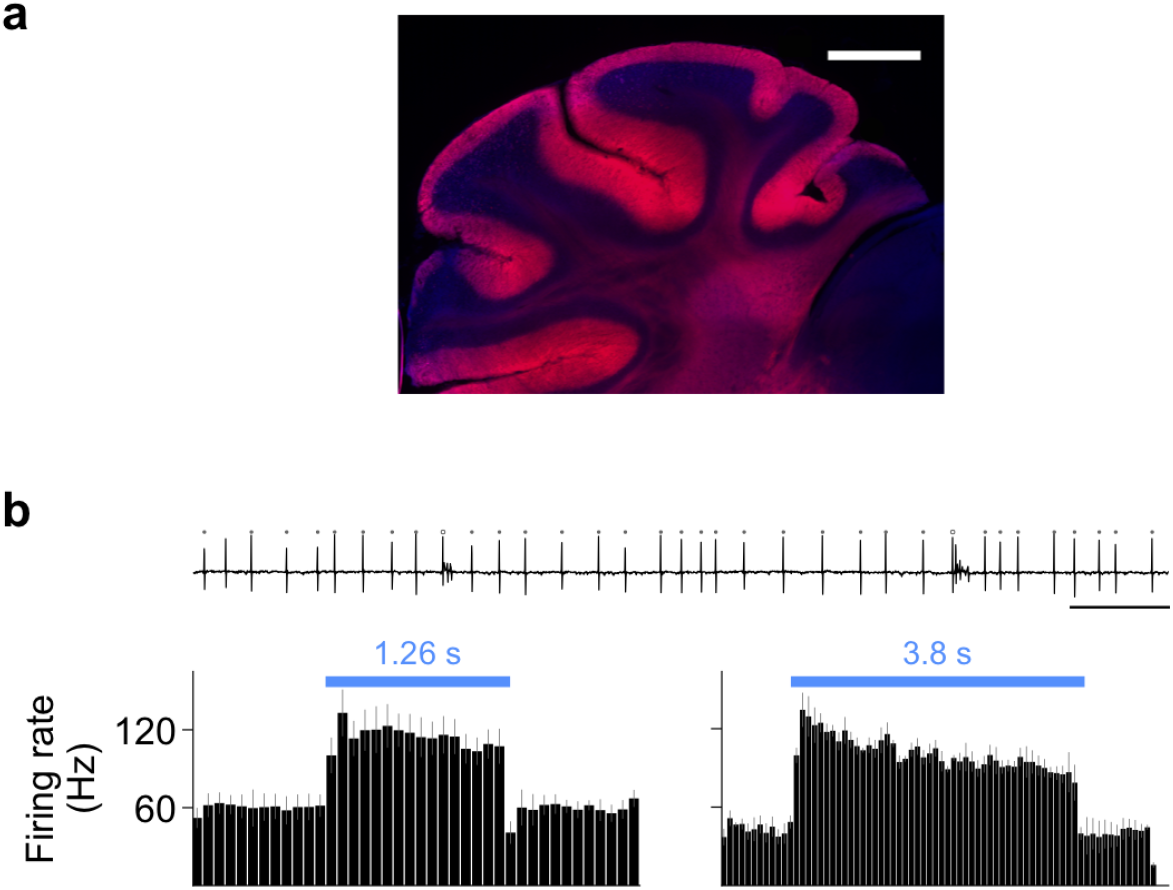
Optogenetic manipulation of cerebellar Purkinje cells. (a) ChR2 expression in cerebellar Purkinje cells. Scale bar: 0.5 mm. Blue: DAPI, red: tdTomato fused to ChR2. (b) Top: extracellular electrical recording from an example Purkinje cell. Circles: simple spikes (filled) and complex spikes (open). Scale bar: 50 ms. Bottom: simple spike firing rate in response to light delivery (blue bars) of different durations. Bars show mean ± s.e.m. of firing rate (n=10 cells, 3 mice). Bin width: 80 ms.

**Supplementary Figure 3:**
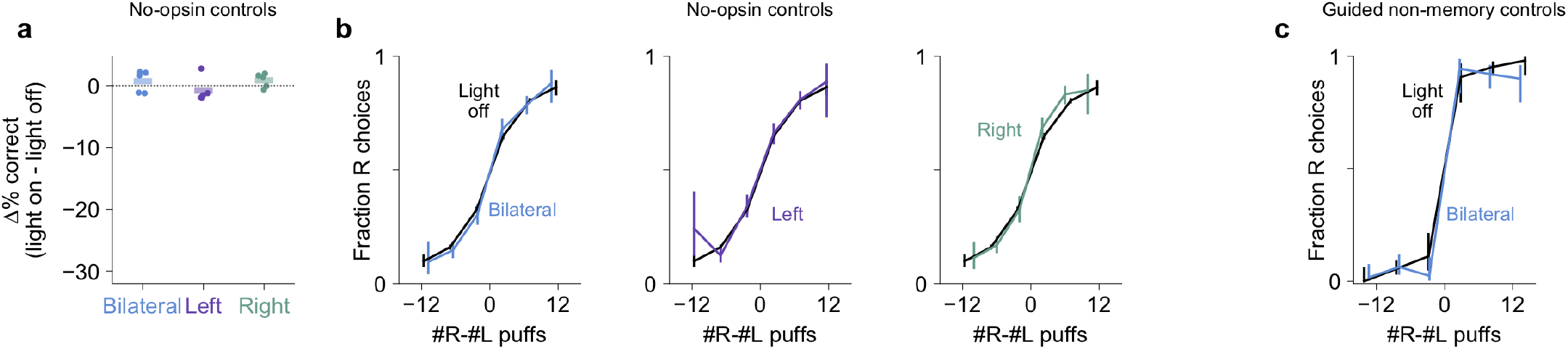
Light delivery has no behavioral effect in ChR2^-^ mice or in non-memory control trials. (a) Change in performance with cue-period light delivery, as in Fig. 1c, for no-opsin control mice (n=15,281 light-off trials, 3,883 light-on trials, 118 sessions, 5 mice). Dots: individual mice. Horizontal lines: mean across mice. Data are not significantly different from zero; left to right: p=0.42, 0.42, 0.17 (two-tailed paired t-tests). Error bars: 95% CI. (b) Psychometric curves as in Fig. 1d for control mice. Error bars: 95% CI. (c) Psychometric curves for experimental mice in trials requiring no memory, where mice were guided to lick the correct side by delivery of all single-sided puffs during the cue period and delay (n=558 light-off trials, 397 bilateral light-on trials, 8 mice). Error bars: 95% CI.

**Supplementary Figure 4:**
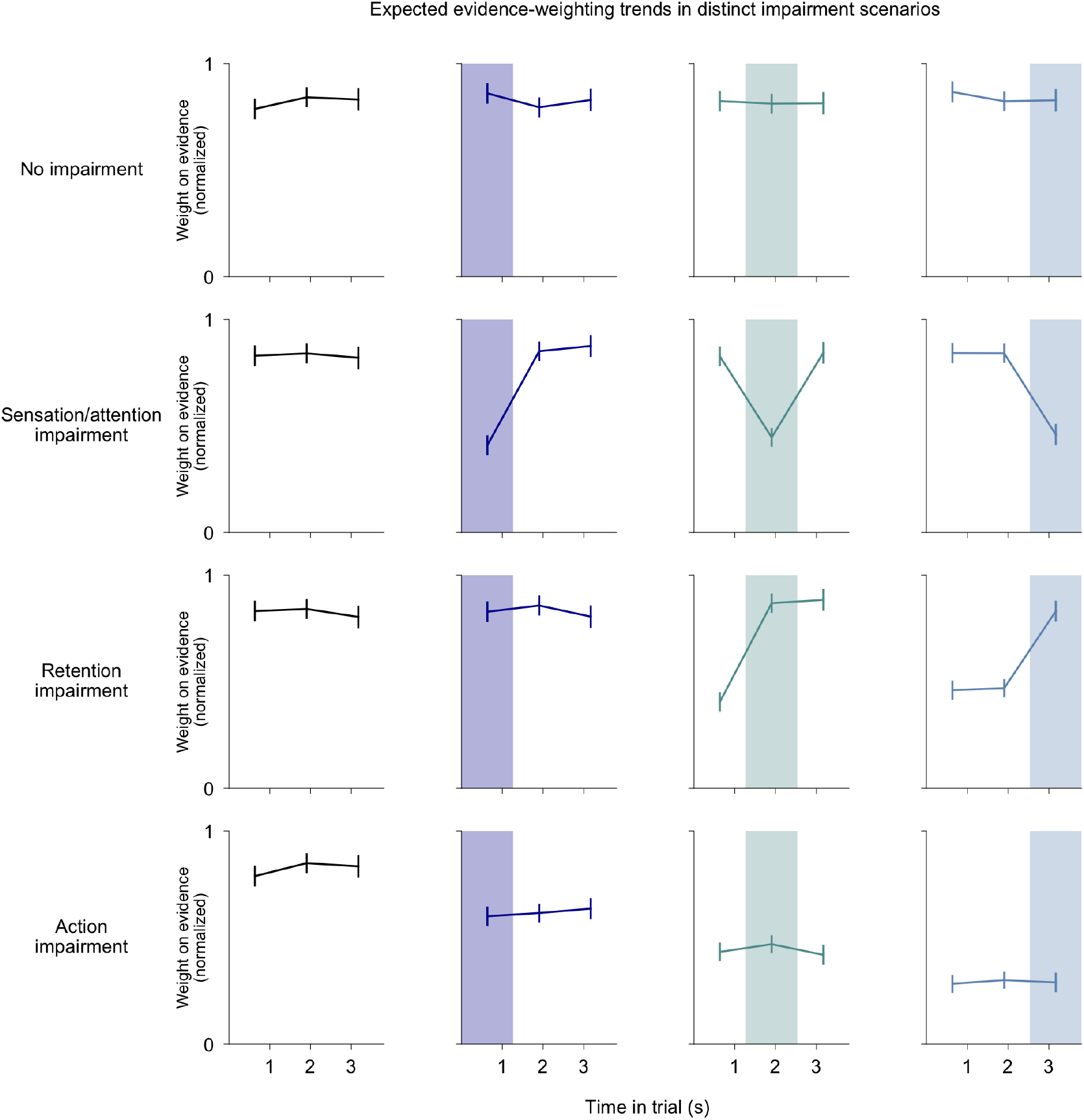
Demonstration of how regression analysis can differentiate distinct perturbation effects. Simulations of different types of perturbations were performed. Trials and associated animal decisions were drawn from the baseline no-perturbation behavioral dataset. Then animal decisions were “perturbed” according to particular rules for four example scenarios (see Methods), presented in the four rows respectively. Each row shows four trial types from left to right, corresponding to light delivery conditions indicated by shading as in Fig. 2 (leftmost column: light off). First row: scenario where light delivery causes no impairment. Second row: scenario where light delivery impairs animals’ ability to sense, encode, or attend to stimuli delivered *concurrently* with the light. Third row: scenario where light delivery impairs the retention of previously accumulated information. Fourth row: scenario where evidence accumulation is intact but light delivery causes a failure to translate the accumulated information into an action, with increasing probability as the decision approaches. All regressions were performed and presented as in Fig. 2a. Regression weights indicate the extent to which evidence was used to guide decisions, and overall performance in any given scenario is proportional to the sum of weights. Error bars: 95% CI.

**Supplementary Figure 5:**
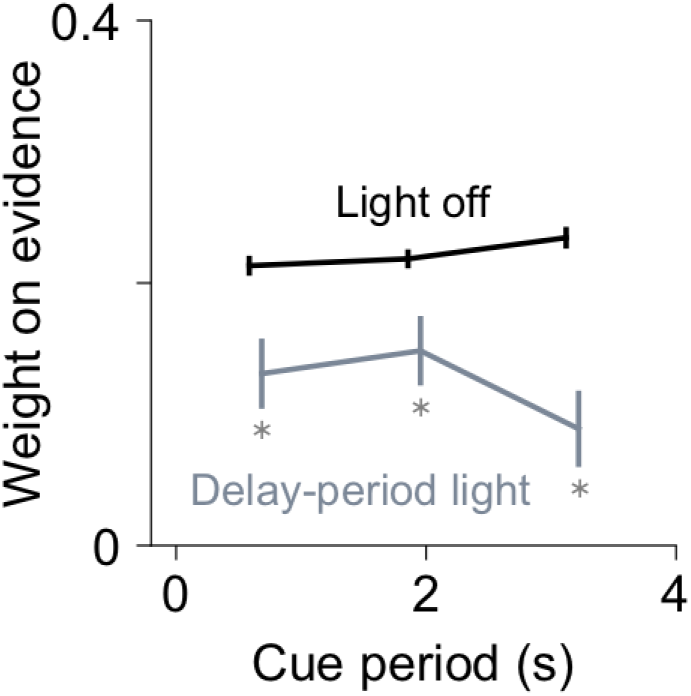
Delay period perturbation. Regression analysis as in Fig. 1e for all light-off (black) and light-on (gray) trials (n=28,959 light-off trials, 2,060 light-on trials, 256 sessions, 8 mice). Weights indicate the extent to which evidence was used to guide decisions, and the sum of weights is proportional to overall performance. *: p<0.05 (95% CI, light-off: 0.2–0.23, 0.2–0.23, 0.22–0.25; light-on: 0.08–0.18, 0.1–0.2, 0.03–0.15).

**Supplementary Figure 6:**
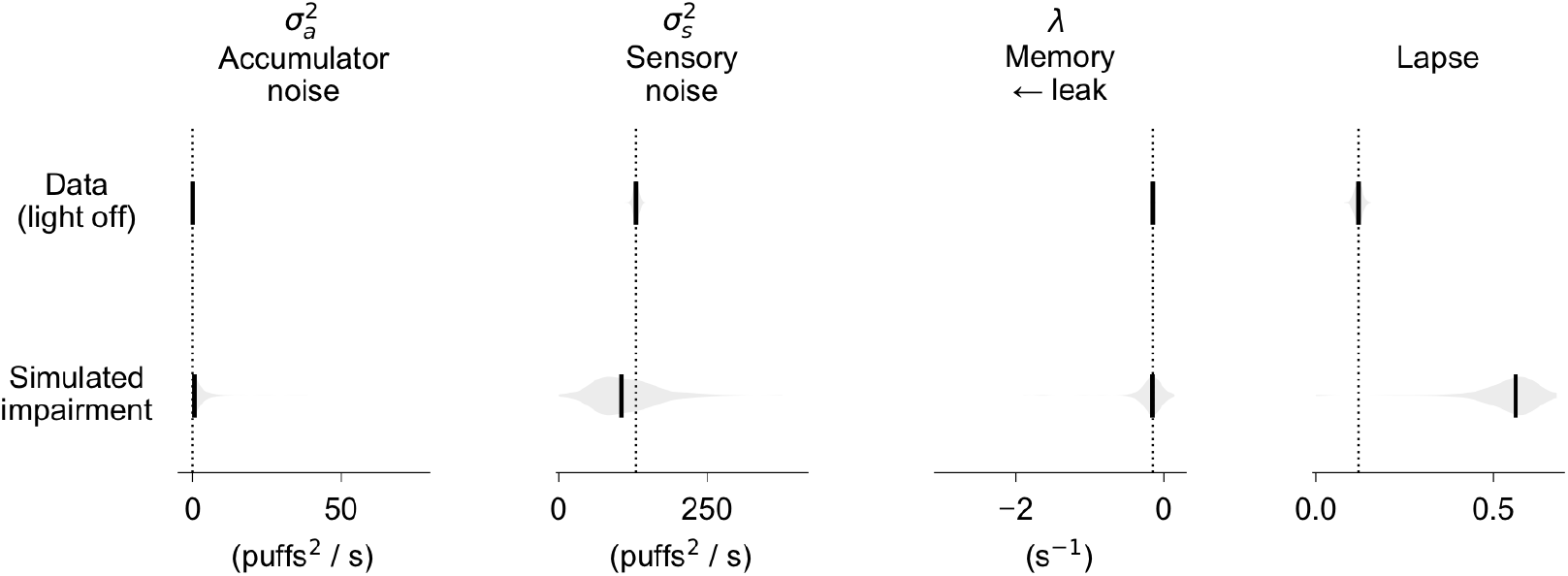
Simulated lapses demonstrate the specificity of model parameters. Subsamples of trials were randomly sampled from the baseline no-perturbation behavioral dataset, and a specific decision impairment was simulated: in a scenario where stimulus accumulation in working memory is intact but animals stochastically fail to translate this information into a directed action, one would observe a random subset of trials in which decisions are opposite of the accumulated information in memory. We therefore simulated the impairment by imposing a random choice on a random subset of trials in these sampled datasets. These trials were then fit to the model in the same manner as the real data (see Methods for additional details). As expected, the model captured the impairment as an increase in lapse rate with no effect on other parameters. This exemplifies the power of the model to identify specific deficits, confirming that the alterations in other parameters with cerebellar perturbation (Fig. 3) are not explained by lapses in animals’ ability to report the information accumulated in working memory. Display conventions are the same as those in Fig. 3.

**Supplementary Movie 1:** Visual demonstration of the drift diffusion model parameters affected by cerebellar perturbation. Four consecutive scenarios are shown, all displaying the same behavioral trial (5 left puffs, 3 right puffs; the correct choice is left). All four scenarios display the model’s accumulator value *a* (i.e. working memory trace of stimulus information) over time as the stimuli are presented. The moving circle and white line display the evolving value of *a*. The color of the circle at any given moment indicates the sign of the accumulator and thus the side the agent would select if the decision occurred at that moment (in the absence of lapses). The arrows indicate puff (evidence) events, and the choice is indicated at the end of the trial by the flashing box. In each scenario, the meaning of one specific parameter is demonstrated.

**Supplementary Table 1:** Best-fit drift diffusion model parameters (95% CI)

| | $\sigma_a^2$<br>(puffs<sup>2</sup> / s) | $\sigma_s^2$<br>(puffs<sup>2</sup> / s) | $\lambda$<br>(s<sup>-1</sup>) | bias | lapse |
| --- | --- | --- | --- | --- | --- |
| Light off | 0.01<br>(0.00 – 0.08) | 129.84<br>(120.62 – 139.08) | -0.15<br>(-0.17 – -0.13) | -0.43<br>(-0.46 – -0.39) | 0.12<br>(0.10 – 0.14) |
| Full cue<br>period light | 33.67<br>(6.72 – 53.63) | 5.54<br>(0.03 – 150.07) | -1.39<br>(-2.77 – -0.24) | 0.48<br>(0.23 – 1.10) | 0.36<br>(0.00 – 0.66) |
| Delay period<br>light | 0.00<br>(0.00 – 1.48) | 84.75<br>(37.30 – 397.57) | 0.11<br>(-0.09 – 0.24) | 1.93<br>(1.37 – 2.55) | 0.48<br>(0.00 – 0.56) |

## Acknowledgements

We thank the members of the laboratories of S.W., D.W.T., Ilana Witten, Carlos Brody, and the BRAIN COGS group for discussion and technical assistance, Marlies Oostland for animal training and discussions, Henk-Jan Boele for comments on the manuscript, and Julia Kuhl for illustration. Funded by National Institutes of Health grants F30 MH115577, U19 NS104648, R01 MH115750, R01 NS045193.

## Author contributions

B.D. performed experiments and analyzed data. M.K. performed histology and electrophysiology experiments. B.D. wrote the manuscript with contributions from all authors. B.D., S.W., and D.W.T. conceived the project.

## Data availability

The datasets generated and analyzed in the current study are available from the corresponding author upon request.

## Code availability

All experimental and analysis code is available at the links provided in the Methods section.

## Methods

### Mice

Experimental procedures were approved by the Princeton University Institutional Animal Care and Use Committee and performed in accordance with the animal welfare guidelines of the National Institutes of Health. Data for the behavioral task came from 13 mice (5 female, 8 male, 8-25 weeks of age during experiments) of genotypes Pcp2-Cre for Purkinje-cell specificity and Ai27D for channelrhodopsin-2 (8 animals *Pcp2*-Cre x Ai27D, 5 animals Ai27D) acquired from The Jackson Laboratory, Stock #010536 (RRID:IMSR_JAX:010536) and #012567 (RRID:IMSR_JAX:012567), respectively. Experimenters were blinded to the genotypes of the mice for the duration of the experiments. Data for electrophysiology experiments came from an additional 3 mice of genotype *Pcp2*-Cre x Ai27D. Mice were housed in a 12-hour: 12-hour reverse light:dark cycle facility, and experiments were performed during the dark cycle. During the experimental day, mice were housed in darkness in an enrichment box containing bedding, houses, wheels (Bio-Serv Fast-Trac K3250/K3251), climbing chains, and play tubes. At other times, mice were housed in cages in the animal facility in groups of 2-4 mice per cage. Mice received 1.0-1.5 mL of water per day. Body weight and condition was monitored daily.

### Surgical procedures

Mice were anesthetized with isoflurane (5% for induction, 1.0–2.5% for maintenance) and underwent surgical procedures lasting 2-4 hours. Two ~500-μm diameter craniotomies were drilled over the cerebellum, one over each hemisphere, directly posterior to the lamboid suture and approximately 3.6 mm lateral to the midline in either direction. Ferrule implants were constructed as in^22^ with 400-μm-diameter optical fiber (Thorlabs FT400EMT) glued to 1.25-mm OD stainless steel ferrules (Precision Fiber Products MM-FER2007-304-4500) using epoxy (Precision Fiber Products PFP 353ND). Ferrules were positioned over each craniotomy with the fiber tip at the surface of the dura mater, and Vetbond (3M) was applied surrounding the exposed fiber. Dental cement (C&B Metabond, Parkell Inc.), darkened by mixing with India ink (Koh-I-Noor #3080-4), was then applied to secure the ferrule to the skull. In some mice, separate implants were placed over neocortex for other experiments. When animals were not engaged in experiments, optical implants were protected using ceramic ferrule sleeves (Precision Fiber Products SM-CS, 1.25-mm ID, 6.6-mm length). Implants were cleaned before each behavior session using a fiber optic cleaning kit (Thorlabs CKF). A custom-machined titanium headplate^23^ was cemented to the skull using dental cement (C&B Metabond, Parkell Inc.). All animals were given buprenorphine (0.1 mg/kg body weight) and rimadyl (5 mg/kg body weight) after surgery and were given at least 5 days of recovery in their home cages before the start of experiments.

### Behavior

Mice were trained to perform a previously described evidence-accumulation decision-making task^17^. Briefly, head-fixed mice were seated in tube for 1-hour behavioral sessions consisting of 200-300 trials. In each trial, independent streams of randomly timed 40-ms air puffs (2.5 Hz, minimum 200 ms interpuff interval) were delivered to the left and right sides over the course of a 3.8-second or 1.5-second cue period (duration chosen randomly with 0.85 and 0.15 probability, respectively). After a delay of 800 ms (or in ~10% of early sessions, 200 ms), lick ports were advanced into the reach of the animal, and animals licked to the side with the greater number of puffs to retrieve a water reward. The animal’s decision was interpreted as the side licked first, regardless of subsequent licks. Guided non-memory trials had the same structure except puffs were delivered only on a single side throughout the cue period, and regular 2.5 Hz guide puffs were delivered during the delay; choice was again defined as the side of the first lick (and in guided trials a reward was delivered in all cases independent of choice). The behavioral apparati were controlled by custom-written Python software (https://github.com/wanglabprinceton/accumulating_puffs).

### Optogenetics

Light for optogenetic stimulation was produced by two 470-nm LEDs (Thorlabs M470F3, one for each implant) each powered by an LED driver (Thorlabs LEDD1B). Fiber optic patch cables (Thorlabs M98L01) carried light from the LEDs to the ferrule implants, where they were connected via custom-machined black delrin sleeves. Light was delivered through 400-μm-diameter optical fibers in 5-ms pulses at 50 Hz (generated by Master-8), with an intensity of 3-15 mW/mm^2^. Based on published results^24–27^, we estimate that the light emitted from each fiber illuminated a roughly spherical region of tissue <1 mm in diameter, corresponding to a large fraction of cerebellar crus I. Light delivery was triggered via electrical signals sent by the behavioral control software through a DAQ card (National Instruments, NI PCI-MIO-16E-4). Cue period light was delivered over the entire cue period through the left, right, or both implants. Sub-cue-period light was delivered bilaterally to both implants for one third of the cue period, and delay period light was delivered bilaterally to both implants for the entire 800-ms delay period or for the first 200 or 500 ms. Light delivery trials were interleaved with light-off trials and were selected randomly with a uniform probability (ranging from 15-30%) throughout the session. All analyses compare light-off and light-on trials only from behavioral sessions in which light was delivered.

### Electrophysiology

Single-unit recordings in 3 awake *Pcp2*-Cre-Ai27D mice were performed using borosilicate glass electrodes (1B100F-4, World Precision Instruments) with 1- to 2-μm tips and 3 to 12 MΩ impedance, fabricated on a pipette puller (P-2000, Sutter Instruments Co.) and filled with sterile saline. Electrical signals were amplified with a CV-7B headstage and Multiclamp 700B amplifier, digitized at 10 KHz with a Digidata 1440A and acquired in pClamp (Axon Instruments, Molecular Devices) in parallel with TTL pulses from a signal generator (Master-8, A.M.P.I.), which was used to synchronize recording and optical stimulation. Light was delivered through a ferrule implant identical to those used in behavior experiments, positioned above an open craniotomy and connected to a fiber-coupled LED (M470F3, Thorlabs) with a TTL-controlled driver (LEDD1B, Thorlabs). The fiber optic was always moved independently of the recording electrode using a second motorized micromanipulator (MP-225; Sutter Instrument Co.). The optical stimulation parameters were the same as those used in the behavioral experiments. Spike detection was performed using custom code written in MATLAB 2017b.

### Histology

Animals were deeply anesthetized and then transcardially perfused using a peristaltic pump with phosphate buffered saline (PBS) followed by chilled 10% formalin (Fisher Scientific). Brains were extracted from the skull after perfusion, postfixed overnight at 4°C, cryoprotected in 30% sucrose in PBS, embedded in O.C.T. compound 4585 (Tissue-Plus, Fisher HealthCare) and stored at -80°C until sectioning. 50-μm-thick sagittal sections were cut with a Leica CM3050 S cryostat. To remove the cryoprotective solution, sections were washed with PBS. Sections were mounted on slides and covered with Fluoroshield anti-fade reagent with DAPI (Sigma). Images were acquired on an inverted fluorescent microscope (Nikon Eclipse Ti) using NIS-Elements AR software. Image processing was performed in Python.

## Data Analysis

### Software

Data analyses and figure creation were performed using custom code written for Python 3.6 (code available at https://github.com/bensondaled/puffsopto), which makes use of Numpy 1. 14.3^28^, Scipy 1.0.0^29^, Pandas 0.23.4^30^, Matplotlib 2.2.2^31^, IPython 6.1.0^32^, Scikit-learn 0.19.1^33^, and Statsmodels 0.9.0^34^.

### Performance and psychometrics

Data for performance and psychometric measures were obtained only from trials in the final stages of the task, and not from the preceding stages during the shaping procedure. Performance, psychometric, and regression analyses contain only trials in which mice made decision licks, such that incorrect trials correspond to licks in the wrong direction, and never the absence of a decision lick. Optogenetic analyses compare light-off and light-on trials only from sessions in which light-on trials were delivered and only from trials with the primary 3.8-second cue period. Confidence intervals on fractions of correct or left/right-choice trials were computed by the Jeffreys method for binomial confidence intervals. The meta-mouse psychometric curve in Supplementary Fig. 1a consists of pooled trials from all mice and was fit to a four-parameter logistic function of the form:

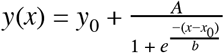

### Behavior regression analysis

To determine the dependence of animal choice on stimuli in different temporal bins of the cue period, we performed a regression-based analysis. Data for regression analysis consisted of trials with a cue period duration of 3.8 seconds. Logistic regressions were performed with animal decision on a trial-by-trial basis as the predicted variable. The input for each trial was a vector of values corresponding to the difference in right vs left puffs in temporally uniform bins of the cue period. Logistic regression models were fit with no intercept term and no regularization. Confidence intervals on regression weights were computed using the standard error of the parameter fits and the standard normal distribution. The light-delivery-aligned regression in Fig. 2c was computed by performing the regression analysis on each perturbation condition separately, then averaging weights across conditions aligned to light onset, wherever these weights existed. For example, the weight following light offset is the mean regression weight at that time point from the first- and middle-third light delivery conditions. Error bars were computed using a bootstrap approach: for each regression fit, a random sample of trials was selected with replacement from the set of trials to be fit, and the analysis was run on these trials. This procedure was repeated 100 times and error bars were computed as the standard deviation of the resulting weights across runs.

### Simulations for regression analyses

For all simulations in Supplementary Fig. 4, we used the full baseline dataset of 48,239 non-manipulation trials delivered to animals during real experiments. In light-off and no-impairment simulations (left column and top row in Supplementary Fig. 4), simulated decisions were sampled trial-by-trial from the empirical psychometric curve exhibited by the trained animals. For light delivery conditions (remainder of panels), the decisions were also simulated in this way, but with the addition of simulated perturbation-like interventions, as follows: (1) in the “sensation/attention impairment” scenario, for each trial, stimuli *coinciding* with light delivery were given half the magnitude of all other stimuli, then the cumulative evidence was summed for the trial yielding a new effective total #R–#L value, from which a decision was drawn using the empirical psychometric curve like above. (2) in the “retention impairment” scenario, for each trial, stimuli *preceding* light delivery were given half the magnitude of all other stimuli, and the same procedure was applied. (3) in the “action impairment” scenario, for each trial, stimuli were summed (i.e. accumulated) normally and decisions were drawn as in the no-impairment condition, but then the decision was stochastically switched to the opposite side with a probability inversely proportional to the time until the decision lick, emulating a failure to execute the decision that matches the agent’s internal accumulated memory. Regressions were performed on each resulting simulation dataset in the same manner as the data figures.

### Drift diffusion modeling

Our model is based on the one presented in ^18^. In each trial, an accumulator value *a(t)* tracks the level of evidence presented in the trial so far, with right-sided stimuli corresponding to positive deflections and left-sided stimuli to negative deflections. When the trial ends, the choice is defined as the sign of *a*, positive for rightward choices and negative for leftward choices. σ_a_^2^ is a diffusion constant that parameterizes noise in *a*. σ_s_^2^ parameterizes noise associated with single left or right puffs. λ parameterizes drift in the memory *a*. When λ < 0, the accumulator *a* drifts towards 0, causing earlier evidence to influence the decision less than later evidence, often called “leakiness.” When λ > 0, the accumulator *a* drifts further from 0, causing earlier puffs to influence the decision more than later puffs, often called “instability.” These features are implemented by the model:

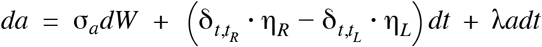

where δ*_t,t_R/L__* are delta functions at the puff events, η are i.i.d. Gaussian variables drawn from *N*(1, σ_*s*_), and *dW* is a white-noise Weiner process. At time *t*=0, the value of *a* is set to 0. In addition, a bias parameterizes an offset in *a* and a lapse rate parameterizes the fraction of trials on which a random response is made. Ideal performance is characterized by an accumulator value *a*=#*R*-#*Lpuffs*, which would be achieved by setting the following parameter values: *λ* = 0, *σ*_a_^2^ = 0, *σ*_s_^2^ = 0, bias=0, lapse=0. Because data were pooled across subjects and bilateral perturbations, we did not interpret the best fit values of the bias parameter to be meaningful, but we included them so as not to capture incidental bias in other parameters like lapse rate. The model was fit using automatic differentiation as in ^7^, and fits were including in analyses only if the resulting Hessian matrix of the model likelihood with respect to the model parameters was positive semidefinite. To estimate the confidence intervals of fit parameters, each model was fit 1000 times, initializing with random values for each parameter and omitting a random 20% of trials in each repetition. The median parameter values and confidence intervals were assessed across fit repetitions.

### Drift diffusion model simulation

The demonstration in the second row of Supplementary Fig. 6 was produced as follows: Random subsamples (n=500 subsamples, 10,000 trials each) were collected from the behavioral dataset without perturbation (i.e. light-off). A simulated “perturbation” was then introduced by choosing a random 50% of trials and replacing the true animal choice with a random selection (either left or right). This reflects the concept of a lapse: i.e. an impairment in selecting the desired response, and specifically one that is not tied to the timing or quantity of accumulated evidence information. Each of the 500 subsamples of trials with the perturbation applied were then fit to the drift diffusion model using the same methods as the data fitting in Fig. 3.

The trials shown in Supplementary Movie 1 and Fig. 3b were generated as follows: a single trial with 5 left puffs and 3 right puffs was produced, and the accumulator value *a* throughout the trial was calculated by running the model (equation in the Drift Diffusion Modeling section above) in discrete time steps of 15 ms. For the Baseline case, parameters were chosen to be similar to the empirically fit light-off behavioral data (Supplementary Table 1). The leaky, noisy, and lapse conditions were simulated by altering those parameters and re-running the simulation. Playback was slowed for visualization purposes.

